# ANTITUMOR SULFATED POLYSACCHARIDES FROM BROWN ALGAE *Dictyota caribaea*

**DOI:** 10.1101/2020.04.04.025320

**Authors:** Alexia Nathália Brígido Assef, Bianca Barros da Costa, Thamyris Almeida Moreira, Luana David do Carmo, Tamiris de Fátima Goebel de Souza, Nylane Maria Nunes Alencar, Ana Paula Negreiros Nunes, Leonardo Paes Cinelli, Diego Veras Wilke

## Abstract

Sulfated polysaccharides (SP) are a complex group of bioactive molecules able to inhibit tumor growth. SP increase the effectiveness of chemotherapy and reduce some side effects. Brown algae produce SP with several biological activities including antitumor. This work aimed to investigate the antitumor effect of SP from the brown algae *Dictyota caribaea* (Dc-SP). Dc-SP were extracted with proteolytic enzyme and supernatant was precipitated with increasing concentrations of ethanol. Antiproliferative activity of Dc-SP was tested by the MTT assay against colon cancer (HCT 116) and metastatic melanoma (B16-F10) cell lines. The antitumor effect was evaluated on Swiss mice transplanted with sarcoma 180 tumor and treated i.p. during 7 days with saline or Dc-SP (25 and 50 mg/kg/animal). Dc-SP did not exhibit cytotoxicity *in vitro*, however the Dc-SP-treated mice depicted up to 50% tumor growth inhibition. Dc-SP treatment induced spleen weight increasing along with intense white pulp disorganization. Furthermore Dc-SP did not depict hepatic toxicity, nephrotoxicity nor leukopenia and did induce increase of platelets count. Altogether, these results represent a promising antitumor host dependent effect induced by Dc-SP.

## Introduction

Cancer is considered one of the most serious diseases, being the main cause of high morbidity and mortality in the world and responsible for more than 8.2 million deaths in recent years (Li, et al., 2017; Li, et al., 2017). Chemotherapy and radiotherapy are widely used as primary treatment in spite of their critical limitations such as restricted efficacy restricted due to multi-drug resistance in tumor cells and its serious adverse effects (Wang et al., 2012). In this way, finding new antitumor drugs with low toxicity and fewer side effects is extremely important. Full filling this need, sulfated polysaccharides (SP) from natural sources depicts antitumor effect with low or lacking cytotoxicity (Choromanska et al., 2015; Li et al, 2019; Barros et al, 2020). Polysaccharides are natural macro-molecular polymers, which can generally be composed of more than 10 monosaccharides of glycosidic bonds in straight or branched chains, with a molecular weight of tens of thousands or even millions (Xie, et al., 2016). Studies have already shown that polysaccharides from different organisms were able to modulate the immune system through the activation of macrophages and the production of inflammatory mediators (Schepetkin et al, 2005; Jin et al, 2008; Yu et al, 2013; Yu et al, 2017). Polysaccharides may have the ability to protect the immune system and some examples already used in the clinic are lentinan and schizophyllan. The synergistic effects of lentinan with various drugs or other therapies have been demonstrated in clinical trials (Neimert-Andersson et al, 2011).

Fucoidans are SP from brown algae which exhibit a range of biological activities, including immunomodulatory, anticoagulant, anticancer, radioprotective and antiviral properties (Ale, Mikkelsen, & Meyer, 2011; Kusaykin et al., 2008). Additionally, as usual for algae polysaccharides, fucoidans have a higher availability, lower production costs and higher recovery rates as advantages for use as functional materials on the industrial scale (Fedorov et al., 2013). Studies have shown fucoidans antitumor effects through cellular apoptosis, inhibiting tumor metastasis and potentiating the toxic effect of chemical drugs (Hyun, et al 2009; Jin, et al 2010; Kim, et al 2010). SP also exert their antitumor effect due to the activation of the immune system of the host animal (Chen et al, 2012; Han et al, 2011). Furthermore, fucoidans usually are non-toxic to organisms and do not have significant side effects. In this study we evaluated the antitumor effect of SP from the brown algae *Dictyota caribaea* (Dc-SP).

## Chemical characterization of sulfated polysaccharides from *D. caribaea* (Dc-SP)

### Material and Methods

#### Sample collection

The marine algae, *Dictyota caribaea*, was collected in March 2011 at Vermelha beach (23°11’46.0”S and 44°38’38.0”W), Paraty, Rio de Janeiro, Brazil, separated of epiphytes and from other species, washed with distilled water, air-dried, powdered and stored at -20°C. Voucher specimens (RFA 38779) were deposited at Herbarium of the Rio de Janeiro Federal University, Brazil (Simas et al, 2014). The use of *D. caribaea* was registered on National System of Management of Genetic Heritage and Associated Traditional Knowledge (SisGen) under the number AC2D331.

#### Sulfated polysaccharides extraction and structure elucidation

##### Proteolytic extraction of SP

Cleaned and dried marine algae (10 g) were immersed in acetone, and kept for 24 h at 4°C. The pellet was dried at 60°C (5 g, dry weight), suspended in extraction buffer (0.1 M NaOAc, 5.0 mM EDTA and 5.0 mM cysteine, 1.0 g of papain in pH 6.0) and incubated at 60°C for 12 h stirring (SL-222, SOLAB, Piracicaba (SP) Brazil) at 200 rpm. The incubation mixture was then centrifuged (LS-3 plus, CELM, São Caetano do Sul (SP), Brazil) (2500 x g for 20 min at room temperature) and the supernatant was saved. The residue was resuspended at the same extraction buffer until absence of SP in supernatant was checked by metachromasia properties with DMB at A 525. Supernatants positive for metachromasia properties were combined, called as crude, and SP were precipitated in the presence of ethanol (at final concentrations of 9%, 23%, 45% and 75%). Each precipitated was dialysed, freeze-dried and stored at -20°C. Approximately 2.7 g (dry weight) of total crude polysaccharides and 1.29 g of F9 were obtained after these procedures (yield of 27% and 12.9%, respectively), calculated using the following equation: yield (m/m, %) = weight of dry seaweed (g) / weight of dry crude polysaccharide (g) x 100. Additionally, the content of sulfated polysaccharides present in F9 was 2.9% calculated using the following equation: yield of sulfated polysaccharide (%) = weight of dry seaweed (g) / biochemical dosage of sulfated polysaccharide (g) x 100.

##### Purification of the SP of seaweed

The SP precipitated in the presence of ethanol was dissolved in 5 mL of 20 mM TRIS-HCl, 50 mM EDTA (pH 7.4), applied to a DEAE-cellulose column (10 cm x 2.0 cm) equilibrated with the same solution and washed with 50 ml of the same buffer. The column was developed by a linear gradient of 0 → 3.0 M NaCl in the same solution. The flow rate of the column was 0.5 mL/min. Fractions of 1.0 mL were collected and assayed for SP using the metachromatic assay with DMB and conductivity. The peak obtained at DEAE-cellulose (Dc-SP) was applied to a High Q – HPLC at the same conditions of equilibration, elution, collection and fraction checked (Cinelli, Vilela-Silva & Mourão, 2009).

###### Animals

A total of 24 Swiss mice (female, 25–30 g), obtained from the central animal house of Federal University of Ceara, Brazil, were used. Animals were housed in cages with free access to food and water. All animals were kept under a 12:12 h light-dark cycle (lights on at 6:00 a.m.). All animal handling procedures were performed in accordance with the Brazilian legislation for the use and care of laboratory animals (No 11.724/2008). This project was approved by the Animal Ethics Committee of the Federal University of Ceara (#NS50).

###### Tumor Cell lines

The cytotoxicity of Dc-SP was tested against the human colon cancer HCT 116 and the murine metastatic melanoma B16-F10 cells. Cells lines were obtained from Banco de Células do Rio de Janeiro, Rio de Janeiro. HCT 116 cells were grown in plastic flasks with RPMI-1640 medium and supplemented with 10% fetal bovine serum, 2 mM glutamine, 100 µg/mL streptomycin and 100 U/mL penicillin and incubated at 37 °C with a 5% CO2 atmosphere. The sarcoma 180 tumor cells were maintained in the peritoneal cavities of the Swiss mice to perform the antitumor assay *in vivo*. Briefly, the ascitic fluid from the abdominal cavity of donor animal was collected and a suspension containing 5 mL of Ringer Lactate, 0.2 mL of Gentamicin (5 mg / mL) and 0.5 mL ascites fluid was prepared. Host animals were inoculated with 2 x 10^6^ cells/0.5 mL intraperitoneally. The procedure was performed every 10 days.

###### Cytotoxic activity – MTT assay

The antiproliferative activity was evaluated by the MTT (3-(4,5-dimethylthiazol-2-yl)-2,5-diphenyltetrazolium bromide) assay (MOSMANN, 1983). The cells were seeded on 96-well plates at a density of 4 × 10^4^ (HCT116 and B16-F10) per well with 200 µL culture medium Briefly, cells were plated 24 hours prior to the addition of the test sample in 96 well plates at the density of 4 x 10^4^ cells/mL. PS was tested in concentration from 1 to 250 µg/mL. Dimethyl sulfoxide (DMSO, 0.05%) was used as negative control and the antineoplastic compound doxorubicin (0.016 to 50 µM) was used as positive control. After treatment, culture media were replaced by fresh media containing MTT solution (0.5 mg/mL) and incubated for additionally 3 h. The MTT solution was, then, removed and the plates dried at 35 °C for 30 min. The formazan product was solubilized in 150 µL of DMSO and the absorbance was read at 570 nm. The inhibition concentration means (IC50) values and their 95% confidence intervals (CI 95%) were calculated by nonlinear regression of the normalized absorbance data to percentage of growth inhibition using GraphPad Prism 6.1 software.

###### Antitumor activity and collection of biological material - Sarcoma 180 tumor model

Eight-day-old sarcoma 180 ascites tumor cells were diluted in lactate ringer solution and transplanted subcutaneously into the left armpit (2×10^6^ cells/500 µl) of the mice. After 24h of the tumor transplant, Dc-SP (25 or 50 mg/kg) dissolved in saline or sterile saline (C-), were admin-istered intraperitoneally for 7 days. At day eight, peripheral blood samples from control and treated mice were collected from the retro-orbital plexus. Then animals were euthanized and tumors, livers, spleens and kidneys were excised, weighed and fixed in 10% formaldehyde. The tumor growth inhibition (%) was calculated by the following formula: inhibition ratio (%) = [(A-B)/A] × 100, where **A** is the tumor weight average of the negative control, and **B** is that of the treated group. Body weights were measured at the day of the tumor transplant and at the last day of treatment. The blood samples were used for hematological and biochemical analyses.

###### Biochemical analysis

Biochemical analyses were performed in serum samples obtained after centrifugation of total blood without anticoagulants, at 2500 rpm for 15 min. Standardized diagnostic kits by LABTEST® spectrophotometer (Lagoa Santa, MG, Brazil) were used in the spectrophotometric determination of alanine aminotransferase (ALT), aspartate aminotransferase (AST) and urea levels.

###### Hematological analysis

For the hematological analysis, an aliquot (∼500uL) of blood per animal was placed in ethylenediaminetetraacetic acid (EDTA) and various hematological parameters (platelet count, total and differential leukocyte count) were carried out by standard manual procedures under light microscopy.

###### Histopathological and morphological analyses

After fixation with formaldehyde, tumors, livers, spleens and kidneys were grossly examined for size or color changes and hemorrhage. Afterwards, tumor, liver, spleen and kidney were cut into small pieces, followed by staining with hematoxylin and eosin of the histological sections. Histological analysis was performed by light microscopy. The presence and extension of liver, kidney or spleen lesions or others possible changes attributed to sample were considered.

###### Statistical analyses

Data are presented as mean and standard deviation (±SD) values. The differences between experimental groups were compared by One-way Analysis of Variance (ANOVA) being followed by Dunnet’s post test to compare weight of the tumor and organs (p<0.05) and Newman-Keuls’ post test to compare the biochemical and hematological parameters (p<0.05) using the GraphPad Prism software v6 (Intuitive Software for Science, San Diego, CA, USA).

## RESULTS

### Dc-SP did not inhibit tumor cell lines growth *in vitro*

The antiproliferative effect of all SP fractions obtained from *D. caribaea* were examined against B16-F10 and HCT116 cells *in vitro*. None of sulfated polysaccharides showed cytotoxic effect until 250 μg/mL (SI.1).

### Dc-SP inhibited tumor growth *in vivo*

The growth of sarcoma 180 was significantly delayed on animals treated with Dc-SP. The tumor mass of the Saline group (treated with 0.9% NaCl) was 2.09 g, while the Dc-SP-treated groups, mean values of 1.15 g and 1.01 g were observed for the 25 and 50 mg/kg/day doses, respectively. The percentages of tumor growth inhibition were 40% (p<0.05) and 51% (p<0.05) for Dc-SP doses of 25 mg/kg and 50 mg/kg compared with negative control (Fig. 1A).

**Figure 1.**
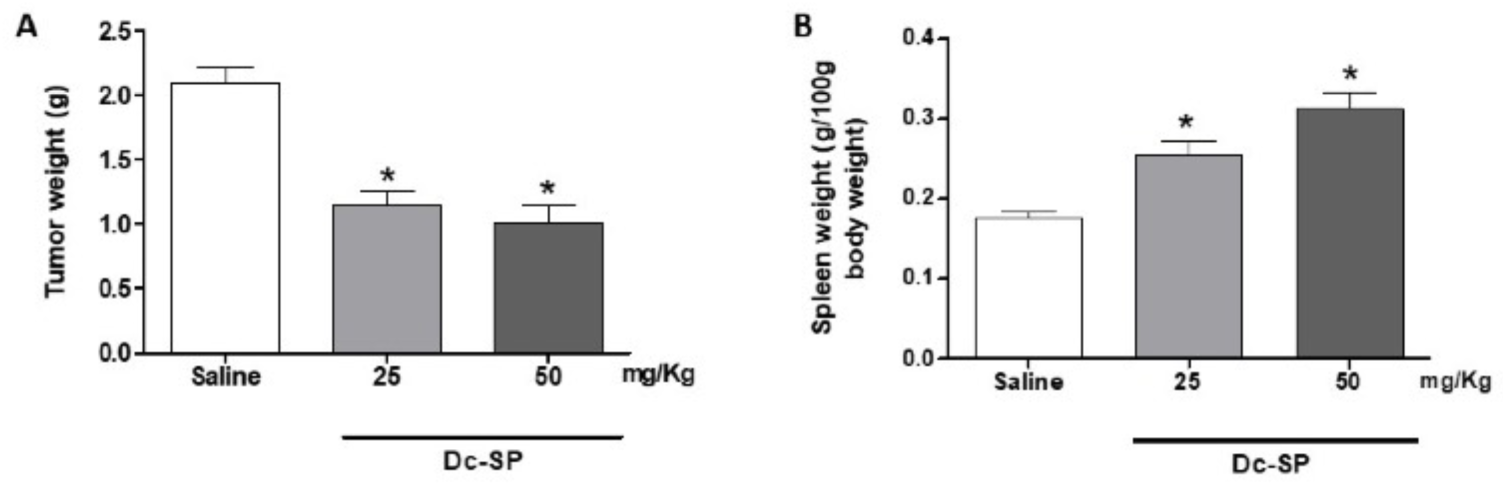
Effects of Dc-SP treatment on the weight of sarcoma 180 tumor (**A**) and spleen (**B**). Mice were treated intraperitoneally with 0.9% NaCl (Saline) as negative control or Dc-SP for 7 days post tumor transplant. Data presented as mean ± standard deviation of N=8 animals per group. *p< 0.05 compared with Saline group analyzed by One-Way ANOVA followed by Dunnet’s test.

### Effect of Dc-SP on organ weight in mice

Dc-SP increased the relative spleen weight of mice dose-dependently in comparison with Saline control group (p<0.05) (Fig. 1B). Dc-SP treatments induced increase of 40% and 70% on spleen weight with 25 and 50 mg/kg doses respectively. The weight of the liver and kidneys of mice were also recorded. There were no change on the weight of the liver (p>0.05) of Dc-SP-treated mice compared with Saline group, however the weight of kidneys increased on Dc-SP-treated mice comparing with Saline group (p<0.05) (SI.2).

### Hematological and biochemical analyses

To determine the effect of Dc-SP on some hematological features of the peripheral blood of mice, comparison was made amongst all the groups. There was a significant (p <0.05) increase in platelet counts in the Dc-SP-treated groups when compared to the saline group (Fig. 2A). The total leukocyte count and the percentage of neutrophils and lymphocytes did not show significant differences in the Dc-SP-treated groups when compared to the saline group (Fig 2B-D). Furthermore differential monocyte counts decreased in the Dc-SP-treated groups when compared to the control group (Fig. 2E). The plasma levels of urea of Dc-SP-treated animals did not change (p>0.05) (Fig. 3A), while the serum levels of aspartate transaminase (ALT) and alanine transaminase (AST) increased when compared to Saline group (Fig. 3B-C).

**Figure 2.**
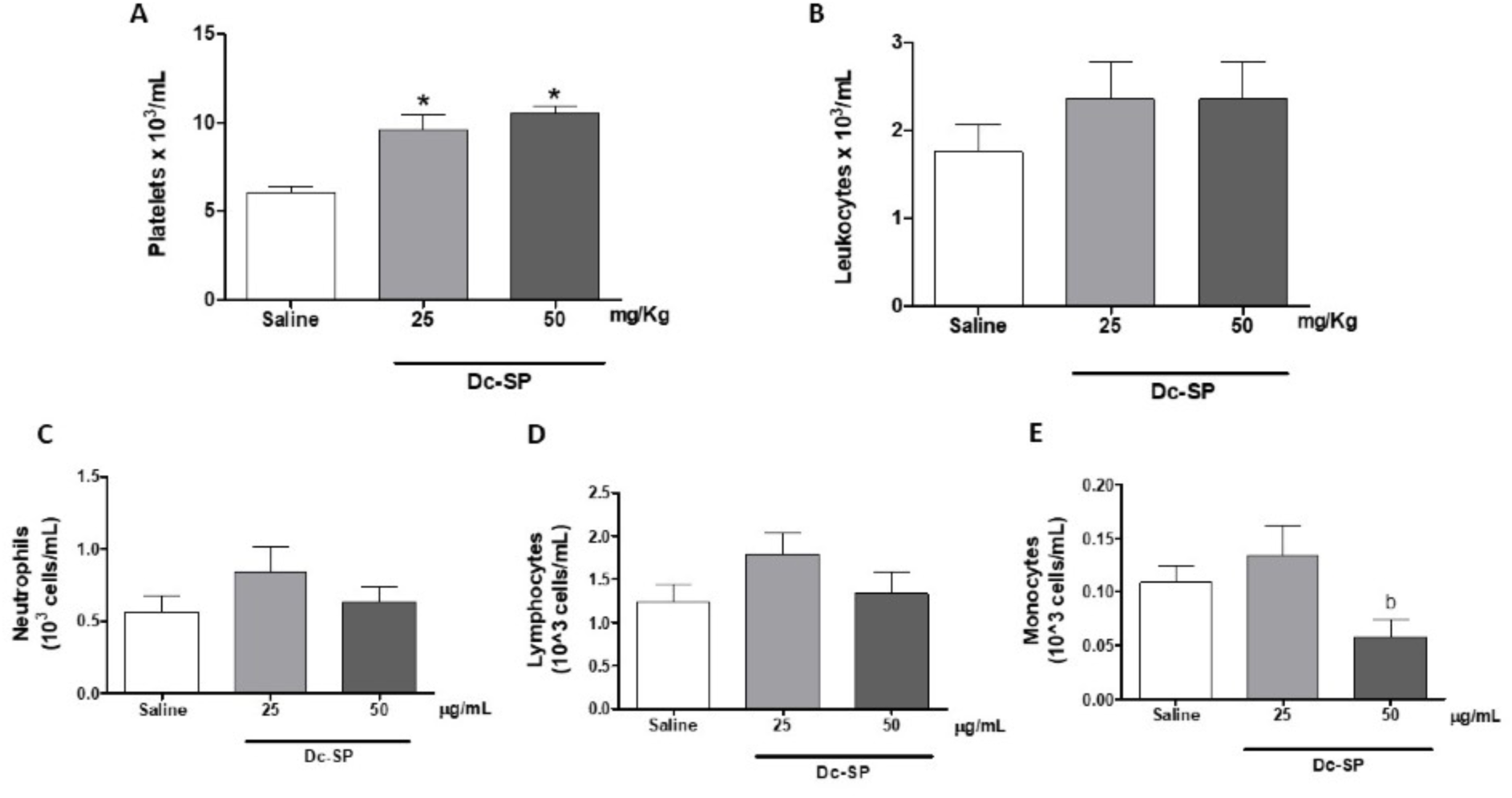
Hematological parameters of peripheral blood from mice transplanted with sarcoma 180. **A**, platelet count, **B**, total leukocytes count, **C**, percentage of neutrophils, **D**, lymphocytes and **E**, monocytes. The animals were treated with 0.9% NaCl (Saline) as negative control or Dc-SP intraperitoneally during seven days, starting from the first day after the tumor implant. Results are expressed as mean ± standard deviation of N=8 animals per group. *p<0.05 compared to Saline group; b compared to Dc-SP 25mg/Kg goup by ANOVA followed by Newman Keuls.

**Figure 3.**
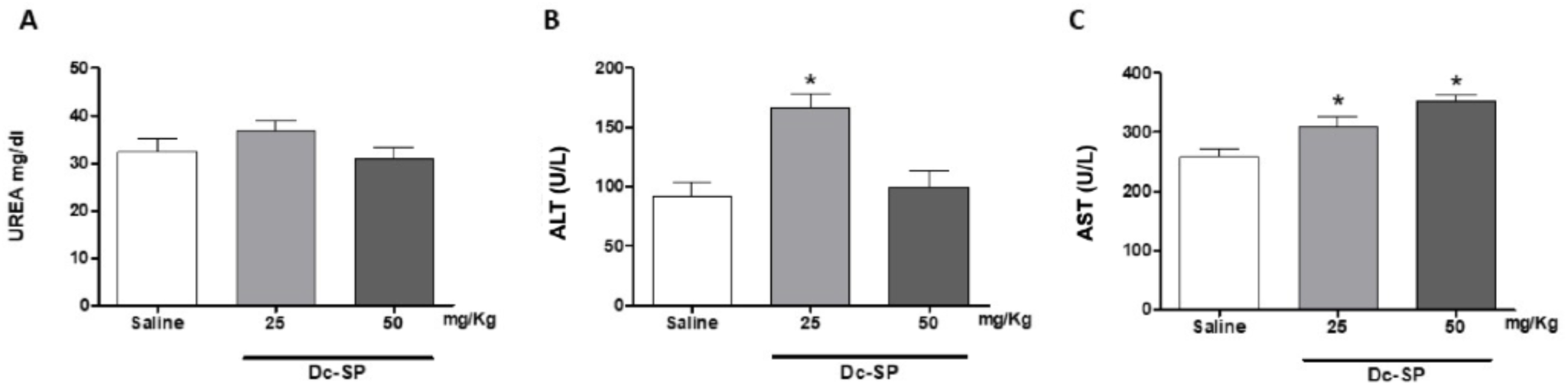
Biochemical parameters of mice transplanted with sarcoma 180. **A**, plasma level of Urea and serum levels of, **B**, aspartate transaminase (ALT) and **C**, alanine transaminase (AST). Animals were treated intraperitoneally for 7 consecutive days from the first day after tumor transplant. The negative control was administered with 0.9% NaCl. Results are expressed as mean ± standard deviation of N=8 animals per group. *p<0.05 compared to Saline group by ANOVA followed by Newman Keuls.

### Histopathological Analysis

The histopathological analysis of the tumors revealed no difference in the morphological pattern among negative control and Dc-SP-treated groups. All groups presented malignant neoplasm with intense cellular disorganization (Fig. 4). Numerous mitotic figures, skeletal muscular invasion and coagulation necrosis were also found on tumors of all groups. Histopathological analyses of the spleen of the animals of all groups showed white pulp hyperplasia, follicle disorganization and presence of megakaryocytes. The spleen of the animals treated with Dc-SP at 25 mg/kg and 50 mg/ kg depicted intense follicular disorganization (Fig. 5). The 50 mg/kg Dc-SP-treated mice also showed congestion of the red pulp. Histopathological analyses of the kidneys of animals treated with saline and Dc-SP showed well preserved renal glomeruli and mild bleeding. The group treated with Dc-SP at 25 mg/kg presented a small vacuolization of the tubular epithelium and necrosis points, whereas the group treated with the dose of 50 mg/kg presented only a small vacuolization of tubular epithelium. All experimental groups presented moderate cellular swelling of the tubular epithelium and glomerular and tubular hemorrhage (SI.3). Histopathological evaluation of the liver of the animals revealed the presence of Kupffer cell hyperplasia and cell swelling in all groups tested. The saline group depicted moderate cellular swelling of hepatocytes. The liver of animals treated with Dc-SP had small inflammatory foci and brown pigments suggestive of bilirubin at both doses (SI.4).

**Figure 4:**
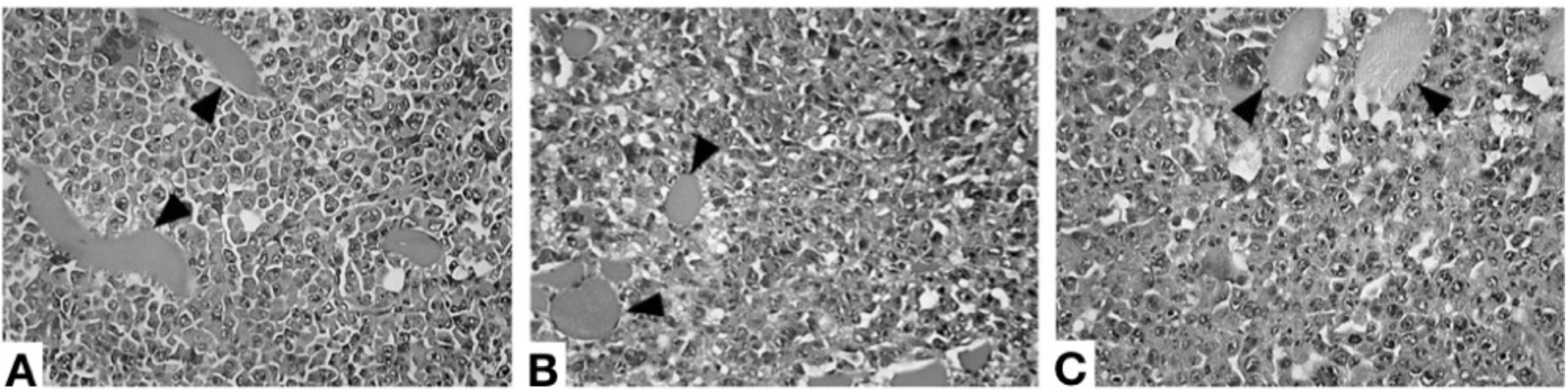
Histological sections of sarcoma 180 tumor. After one day of tumor transplant groups were treated with 0.9% NaCl (Saline) as negative control (**A**), 25 and 50 mg/kg Dc-SP (**B** and **C**) during seven days. Arrows: skeletal muscle fibersevidencing muscle tissue invasion and areas of coagulation necrosis. Hematoxylin and eosin stain. Magnification: 400X.

**Figure 5:**
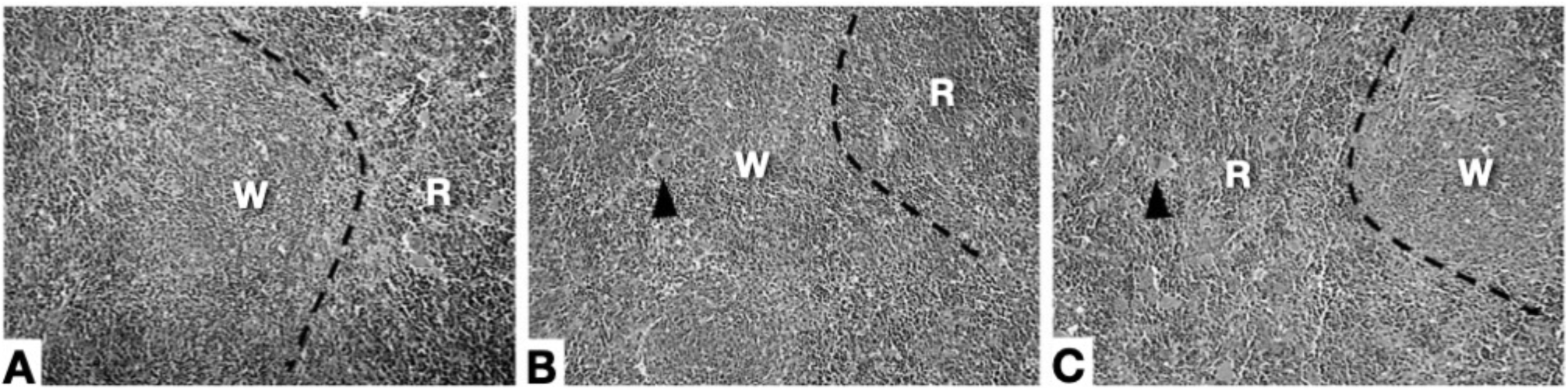
Histological sections of spleen. After one day of tumor transplant groups were treated with 0.9% NaCl (Saline) as negative control (**A**), 25 and 50 mg/kg Dc-SP (**B** and **C**) during seven days. W= white pulp region, R= red pulp region, arrow tips= megakaryocytes. Hematoxylin and eosin stain. Magnification: 400X.

## DISCUSSION

This is the first study showing antitumor activity and immunomodulatory potential of the sulfated polysaccharides from algae *Dictyota caribaea*. Dc-SP treatment inhibited tumor growth and increased spleen weight (Fig. 1), however it did not show cytotoxic activity against tumor cells *in vitro*. Several studies have shown tumor growth inhibition *in vivo* by non-cytotoxic polysaccharides (OOI & LIU, 2000; ZHOU et al., 2004, KHAN et al, 2019; BARROS et al, 2020).

Dc-SP-treated mice increased platelets count (Fig. 2A). Anticancer chemotherapy often causes thrombocytopenia which may lead to haemorrhage and fatality (CIUREA & HOFMANN, 2007). Then Dc-SP effect is desirable and may exert a protective role in patients undergoing such kind of chemotherapy. Polysccharides from the roots of *Angelica sinensis* promote hematopoiesis and thrombopoiesis in mice through PI3K/AKT pathway (LIU et al., 2010). Furthermore Dc-SP-treated mice did not change leukocytes count (Fig. 2B). Leukopenia is also a usual side effect caused by some chemotherapy drugs in clinical use. The use of the chemotherapeutic agent 5-Fluororacil associated with the *Chondrus ocellatus* polysaccharides (ZHOU et al., 2004), *Champia Feldmannii* (LINS et al., 2009) and *Kappaphycus striatum* (YUAN et al., 2006) maintained the antitumor effect and decreased leukocytopenia of 5-Fluorouracil, thus preventing immunodepression promoted by chemotherapy. Additionally the combination of polysaccharides from brown algae (fucoidans) with chemotherapeutic drugs such as cisplatin, tamoxifen and paclitaxel were capable to increase the therapeutic efficacy of the treatments, enhancing apoptosis in cancer cells and showing an increased level of ROS, suggesting the induction of oxidative stress can induce cancer cells death (Zhang et al, 2013).

Dc-SP-treated mice did not depict biologically relevant hepatic or renal toxicities as observed by histopathological and biochemical analyses of these organs. The treatment with *Schinus terebinthifolia* leaf extract produced an increase in AST levels of two to three times. This effect was attributed to the presence of hepatocytes with vacuoles suggestive of microvesicular steatosis (RAMOS et al., 2019). Dc-SP-treated mice increased levels of AST and ALT, however no changes in liver architecture were observed. These results are suitable because the anticancer chemotherapy is very toxic to healthy tissues. The side-effects of chemotherapy are a limiting issue to the chemotherapy success. The association the of Dc-SP with chemotherapy could putatively improve the chemotherapy anti-tumor response without worse of the side effects. However this was not addressed in the present work and should be further investigated.

The Dc-SP antitumor effect was accompanied by increase of the spleen weight of the mice (Fig. 1B). The weight increasing of immune organs is an initial indicator of immunologic activation (Jiang, Zhu & Jiang, 2010). Comparable results were observed in tumor-bearing mice treated with polysaccharides from *Artemisia argyi* (Bao, Yuan, Wang, Liu & Lan, 2013), *Boschniakia rossica* (Wang et al., 2012), *Schisandra chinensis* (Zhao et al., 2013). In addition, the histopathological analysis of spleen of the Dc-SP-treated groups showed intense disorganization of the splenic architecture with hyperplasia of the white pulp and congestion of the red pulp and the presence of several megakaryocytes (Fig. 5). This alteration is compatible with immunostimulatory effect. The proper immune system activation is undoubtedly important to achieve optimal clinical responses of patients under chemotherapy treatment. Polysaccharides from fungi show positive results with chemotherapeutic agents and it is able of prolong administration periods and reduce the adverse effects seen during chemotherapy (ZONG et al, 2012).

Additionally the leukocyte differential count showed decreasing of monocytes in Dc-SP-treated mice when compared to the Saline group (Fig. 2E). The reduction of the number of myeloid cells could be due to cellular activation (MULLER, 2013), the presence of the tumor *per se* (ADAMI et al, 2018) and myelotoxicity (MAJETY et al., 2018). The later did not fit in our results once Dc-SP did not show antiproliferative effects *in vitro* against tumor cells (SI.1) nor macrophages (data not shown). The presence of the tumor *per se* reducing the number of monocytes when compared with health mice (ADAMI et al, 2018). All mice in this study had tumors transplanted, including the Saline group, so the Dc-SP effect on monocytes count was not related to the presence of the tumor. The innate or adaptive immune response depend on the leukocytes crossing the blood vessels by diapedesis. Myeloid cells are key players in cancer either due to anti-tumor and pro-tumor roles depending on the stimuli in the environment (HAAS & OBENAUF, 2019). Then approaches underling shifting pro-tumor features of tumor microenvironment or eliciting immune response are a suitable strategies which reduce the tumor growth. The reduction of monocytes count along with tumor growth delay observed on Dc-SP-treated mice may be related to monocytes activation and deserve further studies. Overall our results clearly showed a promising host dependent antitumor effect induced by Dc-SP.

## 4. CONCLUSIONS

In summary, sulfated polysaccharides from the brown seaweed *Dictyota caribaea* (Dc-SP) delayed tumor growth in mice bearing sarcoma 180 *in vivo* without cytotoxic activity against tumor cells *in vitro*. The host dependent antitumor effect could be related to an immunostimulant effect. Furthermore Dc-SP treated mice did not show toxicity. These findings highlight a promising bioactive profile of Dc-SP. Further investigations on the elucidation of the mechanisms of anticancer and immunomodulatory effects will guide the potential use of DC-SP on chemotherapy treatment.

## Supplementary Information

**SI.1.**
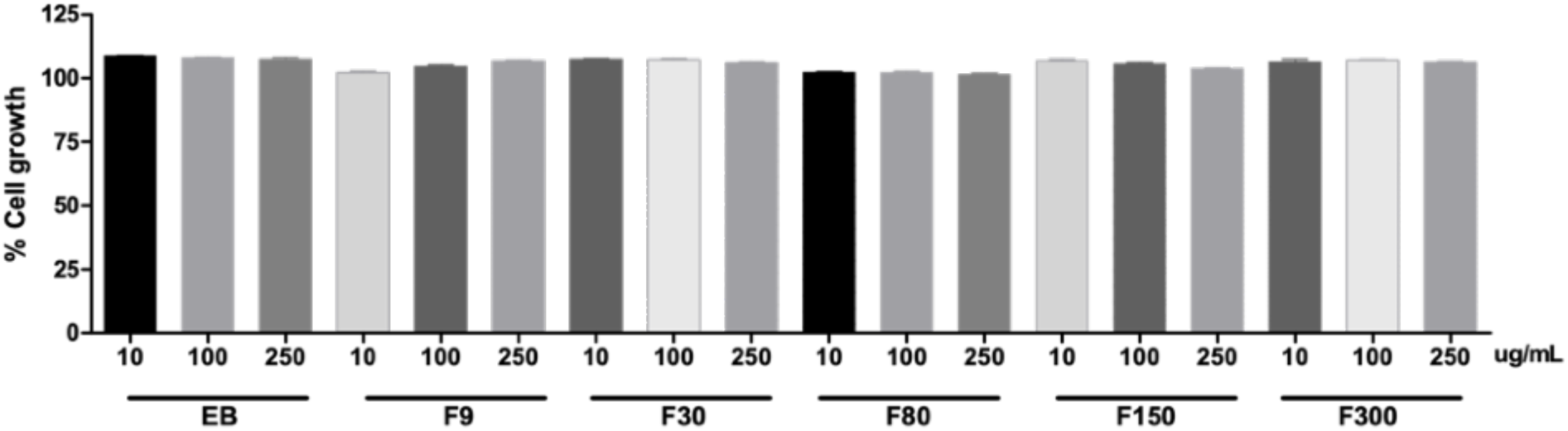
Cell growth of melanoma B16-F10 of treatment with 10, 100 and 250 µg / mL of the PS from *Dictyota caribaea* by MTT assay after 48h. EB = crude extract, F9-F300 = SP fractions.

**SI.2.**
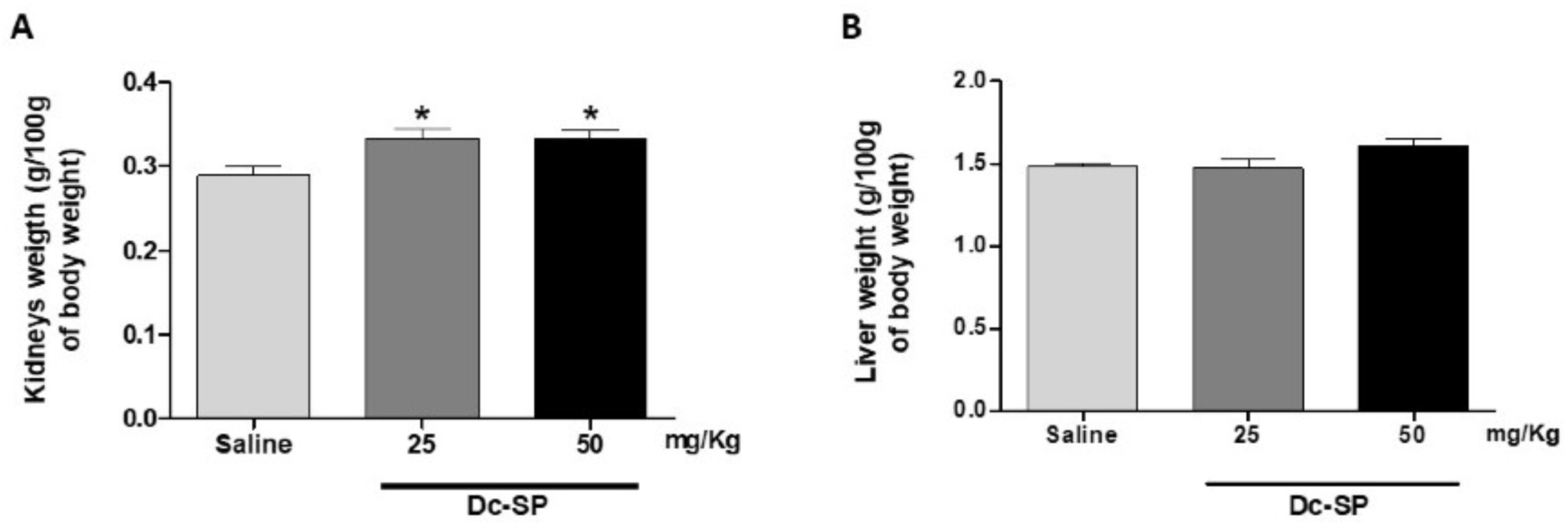
Kidneys and liver weight in mice treated with Saline (C-) and Dc-SP 25 or 50 mg/Kg for 7 days after transplant of Sarcoma 180. *p< 0.05 compared with saline group analyzed by One-Way ANOVA followed by Dunnet’s test.

**SI.3.**
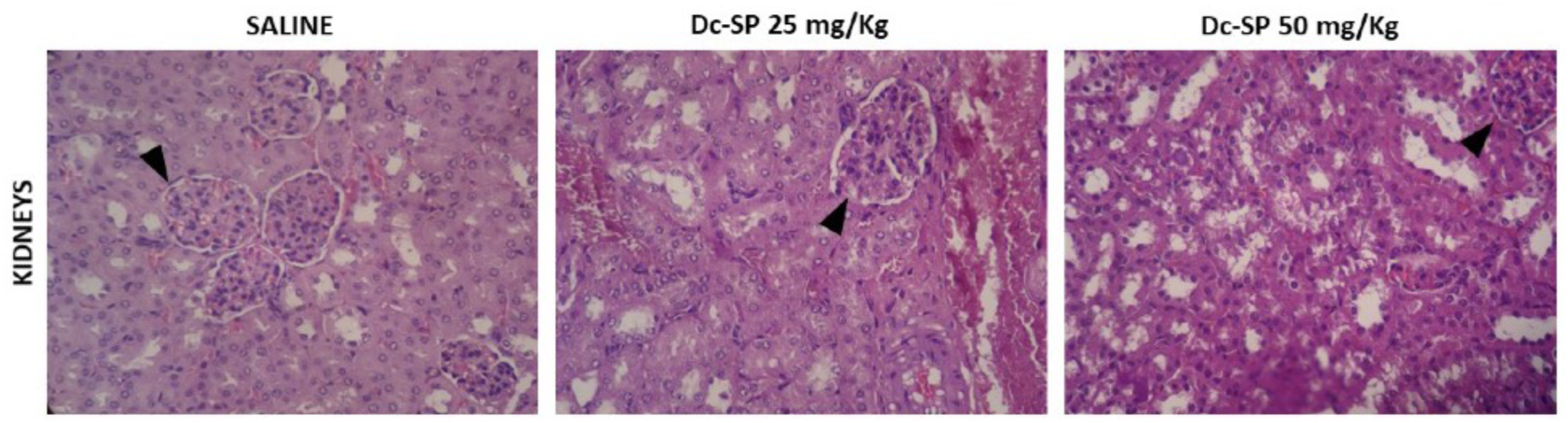
Photomicrographies of kidneys showing well preserved renal glomeruli (arrow tips) and mild bleeding. Groups treated with Saline or Dc-SP at 25 and 50 mg/kg/animal/day during seven days. Histological sections were stained with hematoxylin and eosin. Magnification: 400X.

**SI.4.**
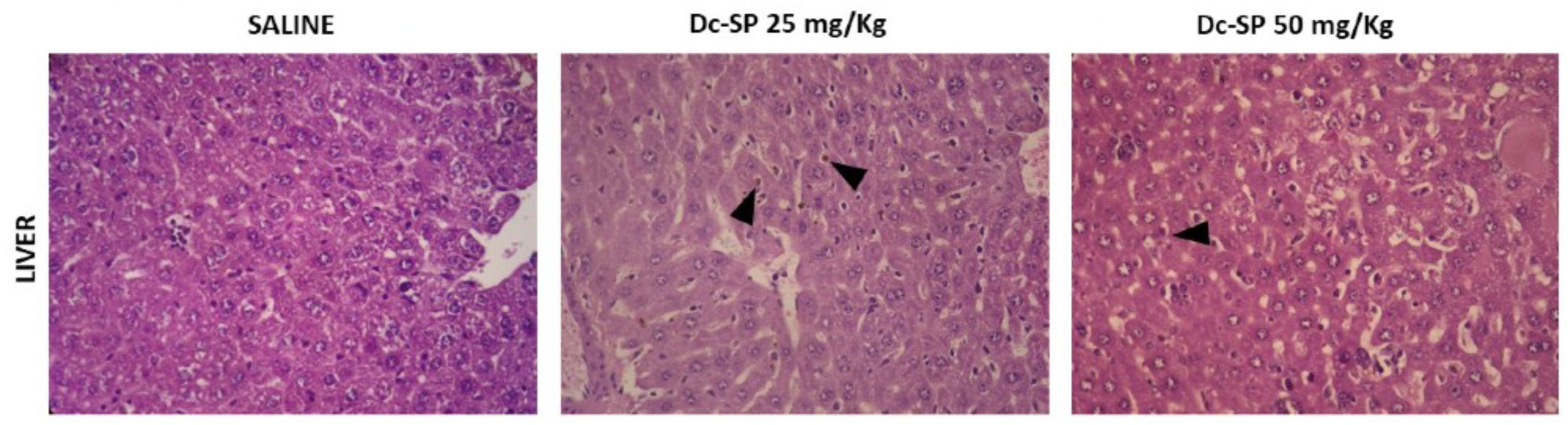
Photomicrographies of Livers. Inflammatory foci and brown pigments suggestive of bilirubin (arrow tips). Groups treated with Saline or Dc-SP at 25 and 50 mg/kg/animal/day during seven days. Histological sections were stained with hematoxylin and eosin. Magnification: 400X.

